# AncestryGrapher toolkit: Python command-line pipelines to visualize global– and local– ancestry inferences from the RFMix2 software

**DOI:** 10.1101/2023.12.29.573635

**Authors:** Alessandro Lisi, Michael C. Campbell

**Affiliations:** Department of Biological Sciences (Human and Evolutionary Biology Section), University of Southern California, 3616 Trousdale Parkway, Los Angeles, CA 90089

**Keywords:** genetic ancestry, admixture, chromosome painting, population genetics, lactase persistence

## Abstract

**Summary:** Admixture is a fundamental process that has shaped patterns of genetic variation and the risk for disease in human populations. Here, we introduce the AncestryGrapher toolkit for visualizing inferred global– and local– ancestry by the `RFMix v.2` software. Currently, there is no straightforward method to summarize population ancestry results from `RFMix` analysis.

**Results:** To demonstrate the utility of our method, we applied the AncestryGrapher toolkit to the output files of `RFMix` v.2 to visualize the global and local ancestry of individuals in the Mozabite Berber population from North Africa. Our results showed that the Mozabite Berbers derived their ancestry from the Middle East, Europe, and sub-Saharan Africa (global ancestry). Furthermore, we found that the population origin of ancestry varied considerably along chromosomes. More specifically, we observed variance in ancestry along chromosome 2 (local ancestry), in the genomic region containing the common genetic polymorphisms associated with lactase persistence, a trait known to be under strong positive selection. This finding indicates that the demographic process of admixture has influenced patterns of allelic variation in addition to natural selection. Overall, the AncestryGrapher toolkit facilitates the exploration, interpretation, and reporting of ancestry patterns in human populations.

**Availability and implementation:** The AncestryGrapher toolkit is free and open source on https://github.com/alisi1989/RFmix2-Pipeline-to-plot.

## 1 INTRODUCTION

Admixture, defined as gene flow between previously diverged source populations leading to new populations with ancestry from multiple sources, has been an important contributor to levels and patterns of genetic variations in humans (Rudan 2006; The 1000 Genomes Project Consortium 2012; Hellenthal *et al*. 2014; Gurdasani *et al*. 2015; Uren *et al*. 2016; Korunes and Goldberg 2021). For example, a number of studies have shown that regions of the modern human genome have originated from archaic hominins, such as Neanderthals and Denisova (Qin and Stoneking 2015; Sánchez-Quinto and Lalueza-Fox 2015; Sankararaman *et al*. 2016; Wall and Yoshihara Caldeira Brandt 2016; Browning *et al*. 2018; Vyas and Mulligan 2019; Koller *et al*. 2022; Vespasiani *et al*. 2022; Witt *et al*. 2023). Other analyses have also revealed that admixture among modern human populations has occurred over millennia, leading to complex patterns of genomic diversity (Campbell and Tishkoff 2008; Campbell *et al*. 2014; Bekada *et al*. 2015; Arauna *et al*. 2016; Korunes and Goldberg 2021; Gopalan *et al*. 2022).

To characterize patterns of admixture, several computational methods have been developed to infer global ancestry among individuals (Pritchard, Stephens and Donnelly 2000; Alexander, Novembre and Lange 2009) and local ancestry along chromosomes within individuals (Tang *et al*. 2006; Sankararaman *et al*. 2008; Sundquist *et al*. 2008; Price *et al*. 2009; Maples *et al*. 2013; Durand *et al*. 2014; Guan 2014; Dias-Alves, Mairal and Blum 2018; Montserrat, Bustamante and Ioannidis 2020). Of these methods, `RFMix v.1.5.4` is considered to be state-of-the-art in estimating ancestry in complex admixture scenarios (Daya *et al*. 2014; Uren, Hoal and Möller 2020; Hilmarsson *et al*. 2021). Furthermore, an updated version of this software—called `RFMix v.2`—was recently developed and is argued to be superior to the previous version (Carrot-Zhang *et al*. 2021). However, there is currently no straightforward approach to visualize the global and local ancestry results from `RFMix v.2`, which is critical for the exploration, interpretation, and reporting of ancestry patterns in human populations. To address this challenge, we developed computational tools to visualize both global and local ancestry in individuals from admixed populations inferred by the `RFMix` v. 2. software.

## 2 IMPLEMENTATION

Inferences of genetic ancestry are informative for mapping the population origins of genetic risk alleles associated with complex diseases (Freedman *et al*. 2006; Cheng *et al*. 2009; Daya *et al*. 2014) and for understanding the genetic history of admixed populations, including the population origins of admixture events (Daya *et al*. 2014; Uren, Hoal and Möller 2020; Browning, Waples and Browning 2023).

The AncestryGrapher toolkit enables users to visualize global and local ancestry with two distinct pipelines, Global Ancestry Painting (GAP) and Local Ancestry Painting (LAP; Figure 1), that run in a command-line Terminal window on Mac OS X and Linux machines. To execute these pipelines on a Microsoft Windows computer, users will need to install the Anaconda command-line environment on their host machine.

**Figure 1:**
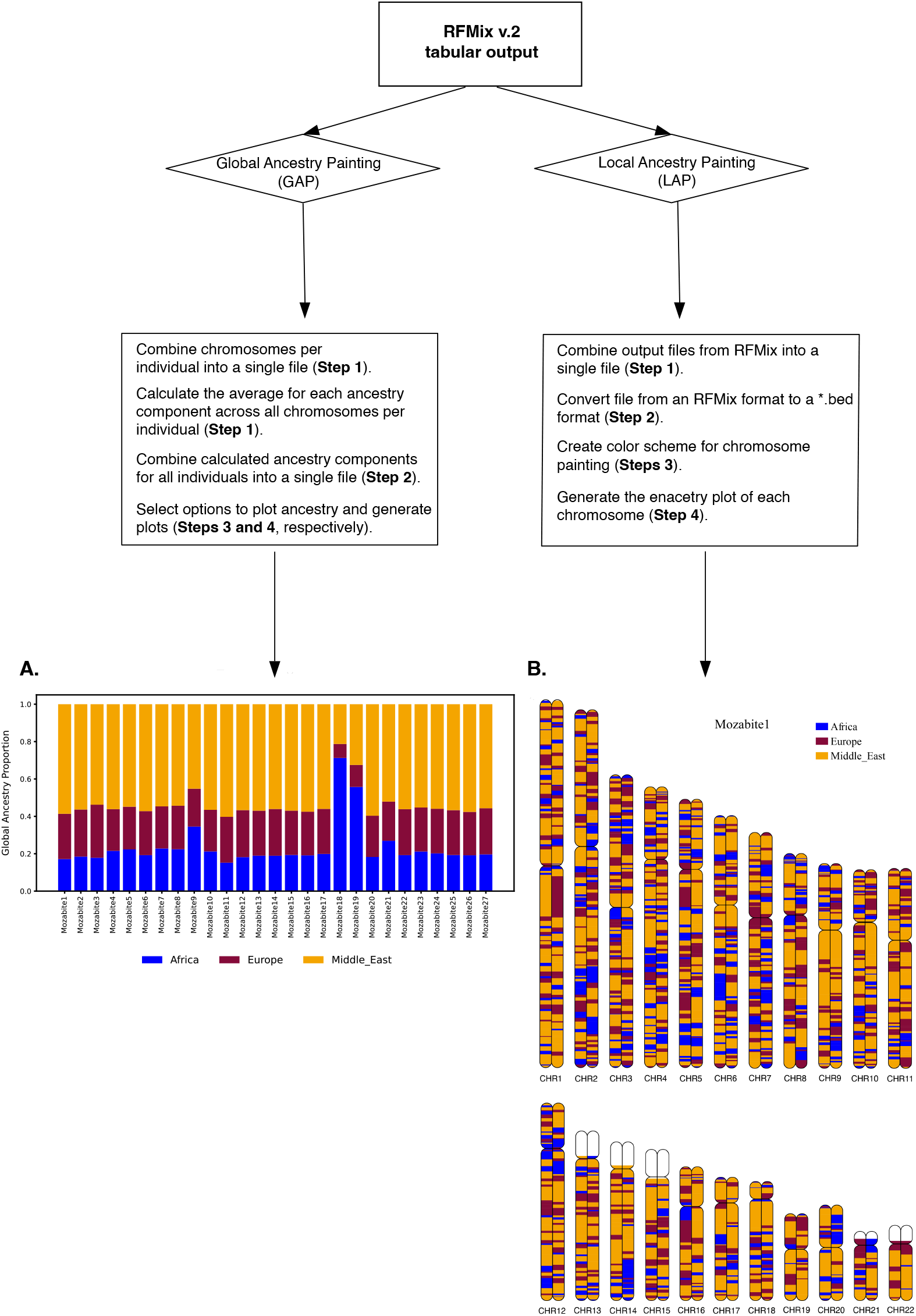
Workflow of the AncestryGrapher toolkit. AncestryGrapher consists of two pipelines: 1) Global Ancestry Painting (GAP) and 2) Local Ancestry Painting (LAP) that run in a command-line Terminal. GAP combines chromosomes into a single file, calculates the average ancestry across chromosomes per individual, and visualizes their ancestry in a bar plot (Panel A). LAP combines the output from `RFMix v.2` into a single file, creates a color scheme for ancestry painting, and plots this ancestry along chromosomes (Panel B). The orange color indicates mean Middle Eastern ancestry; purple represents mean European ancestry; blue signifies mean sub-Saharan African ancestry; the white color represents regions of the genome with no available SNP data.

The AncestryGrapher toolkit can be downloaded to users’ local computers by pressing the “code” button shaded in green on the Github page (https://github.com/alisi1989/RFmix2-Pipeline-to-plot.git). Using the command-line interface in the Terminal window, users will change the current working directory to the directory where the downloaded “RFmix2-Pipeline-to-plot-main.zip” folder is located. To unzip this folder, type “unzip RFmix2-Pipeline-to-plot-main.zip” at the command line prompt (typically indicated by a “$” sign) and the uncompressed “RFmix2-Pipeline-to-plot-main” folder will appear. To demonstrate the utility of our method, we applied the AncestryGrapher toolkit to the output files from an `RFMix v.2` analysis of the Mozabite Berber population from North Africa.

### 2.1 Global Ancestry Painting (GAP)

The GAP pipeline consists of four separate Python scripts: 1) individuals_collapse.py; 2) RFMix2ToBed4GAP.py; 3) BedToGAP.py (this script creates the input file for GAP); 4) GAP.py. Users will need to change their working directory to “GlobalAncestryPaint” located inside of the “RFmix2-Pipeline-to-plot-main” directory.

#### Step 1: Combine the output files for all the chromosomes per individual into a single file

The `RFMix v.2` software generates a *.rfmix.Q (Global ancestry information) output file for each chromosome per individual. Thus, there are multiple files for each individual in a given dataset. The individuals_collapse.py Python script will combine chromosomes for each individual into a single file and then calculate the average of each ancestry component across chromosomes. For example, if there are ten individuals in a dataset, this script will generate ten separate output files with a mean value for each ancestry component for a given individual. Alternatively, if users wish to analyze a single chromosome only, they can proceed directly to Step 2.

#### Step 2: Merge all the individuals or all the chromosomes into one file

To combine the individual files from Step 1 into a single file that contains all individuals under study, users will run RFMix2ToBed4GAP.py.

#### Step 3: Create the input file for GAP to visualize the global ancestry proportions

The output file generated in Step 2 will serve as the input file for BedToGAP.py. Here, there are additional options that users should consider. Specifically, users can incorporate up to five distinct colors, one for each ancestry component with the --ancestry flag. The --ancestry flag accepts two arguments: population ancestry name and the hex color code, which can be found on any website on the internet (*e*.*g*., computerhope.com).

#### Step 4: Generate the plot for global ancestry proportion

In this step, users will run the GAP.py script to generate an output file with ancestry proportions for all individuals in a bar plot together with a legend of ancestry origin in either “pdf” or “svg” format (Figure 1).

### 2.2 Local Ancestry Painting (LAP)

The LAP pipeline consists of three separate Python scripts: 1) RFMIX2ToBed.py; 2) BedToLAP.py; 3) LAP.py.

#### Step 1: Combine the output files from RFMix v.2 into a single file

`RFMix v.2` generates a *.msp.tsv (Local ancestry information) output file for each chromosome for a given individual. Users will combine these chromosomes into a single file, which will contain header and ancestry information for all chromosomes for each individual in the dataset.

#### Step 2: Convert RFMix v.2 output files to *.bed files

In this step, the output file from step 1 will serve as the input file in Step 2. We recommend that users acquaint themselves with the usage of the RFMix2ToBed.py script by typing “python RFMix2ToBed.py -- help” at the command line prompt (typically indicated by a “$” sign). This script will generate two *.bed output files.

#### Step 3: Create the color scheme for ancestry painting along chromosomes

The two *.bed output files from Step 2 will serve as the input files for the BedToLAP.py script. Here, users can enter up to five distinct colors (color codes), one for each ancestry component using the –ancestry flag. This flag requires a single argument, specifically a user-defined hex color code. If no color is specified, default colors will be selected. Users also have the option of highlighting specific genes or genomic regions on a chromosome by inputting additional parameters.

#### Step 4: Generate the ancestry plot for each chromosome for a given individual

The resulting output files from LAP.py will contain ancestry-informative karyotypes along with a legend of ancestry origin and the name of the individual. In addition, the images in the output files will have 4k resolution (4210 x 1663) and can be edited in Adobe Illustrator or Inkscape.

## 3 RESULTS AND DISCUSSION

We have developed the AncestryGrapher toolkit to visually summarize the output of the `RFMix v.2` software. To demonstrate the utility of our computational pipelines, we applied the GAP and LAP algorithms to plot the inferred ancestry of individuals in the Mozabite Berber population determined by `RFMix v.2`. These results showed that individuals in this population have distinct levels of Middle Eastern, European, and sub-Saharan African ancestry (Figure S1; Figure S2), which are consistent with prior studies that argued contemporary North African Berbers are the descendants of the earliest back-to-Africa migration events from the Middle East followed by admixture with populations from neighboring geographic regions (Henn *et al*. 2012; Bekada *et al*. 2015; Arauna *et al*. 2016; Silva *et al*. 2021). In the end, our computational pipeline allows for the easy comparison of global ancestry amongst individuals which cannot be achieved using the standard output of the `RFMix v.2` software (Table 1 and Table S1).

**Table 1.**
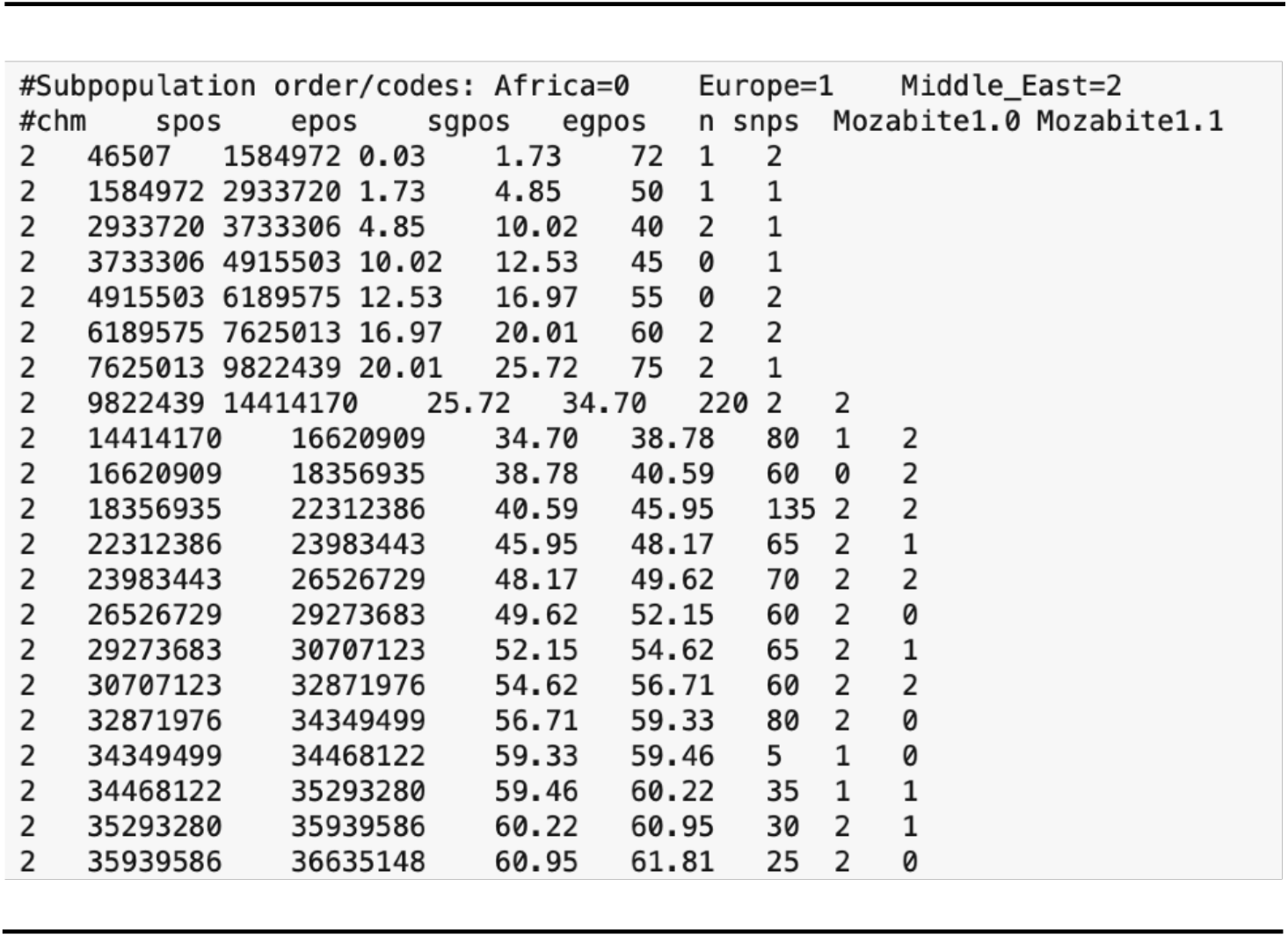
Output file containing local ancestry information for a Mozabite Berber individual generated by RFMix v.2. The local ancestry output files are tab-separated values that correspond to the following: chromosome number (column 1); start and end positions in base pairs (column 2 and column 3, respectively); start and end genetic map positions (columns 4 and 5, respectively); the number of single nucleotide polymorphisms (SNPs) within a given window; and the population ancestry within a given window for each of the diploid chromosomes. The order and names of the reference populations used in ancestry determination are given in the header row.

We also observed striking patterns of local ancestry along individual chromosomes. For this analysis, we focused our attention on the well-studied *LCT* region, which contains several common single nucleotide polymorphisms—namely, C/T_−13910_ (rs4988235), C/G_−13907_ (rs41525747), T/G_−13915_ (rs41380347), T/G_−14009_ (rs869051967), and G/C_−14010_ (rs145946881)—associated with the lactase persistence (LP) trait in human populations (Enattah *et al*. 2002; Ingram *et al*. 2007, 2009; Tishkoff *et al*. 2007; Hassan *et al*. 2016; Anguita-Ruiz, Aguilera and Gil 2020; Campbell and Ranciaro 2021). Prior studies have also shown that these functional alleles have been maintained in Middle Eastern, European, and African populations due to the action of recent selection (Bersaglieri *et al*. 2004; Tishkoff *et al*. 2007; Enattah *et al*. 2008; Gerbault *et al*. 2009; Itan *et al*. 2009; Jones *et al*. 2013; Macholdt *et al*. 2014; Ranciaro *et al*. 2014; Scheinfeldt *et al*. 2019; Vicente *et al*. 2019; Hollfelder *et al*. 2020).

Based on our plots, we found that the LP-associated variants in the Mozabite Berber population sit on chromosomal segments (on chromosome 2) originating from the Middle East, Europe, and/or sub-Saharan Africa. More specifically, the *LCT* region was found to sit in a chromosomal region originating from the Middle East on 46.3% of chromosomes, while the *LCT* region had ancestry backgrounds originating from Europe and sub-Saharan Africa on 40.7% and 13% of chromosomes, respectively, in the Mozabite Berbers (Figure S3). Thus, a high proportion of the genetic variation associated with LP in the Mozabite Berbers was likely introduced into this population through admixture, which we visually captured using the LAP pipeline.

In summary, many human populations have experienced admixture throughout their history with extensive examples of this process in Africa (Campbell and Tishkoff 2010; Campbell *et al*. 2014). Thus, a natural extension of the `RFMix` tool is to visually represent these genetically complex populations to assist with the exploration, interpretation, and reporting of results that could be informative for anthropological, biological, and biomedical research.

## AUTHOR CONTRIBUTIONS STATEMENT

A.L.: Conceptualization, Methodology, Software, Formal Analysis, and Visualization.

M.C.C.: Conceptualization, Methodology, Resources, Writing - Review & Editing, Supervision, Project administration, Funding acquisition.

## ACKNOWLEDGEMENTS

We thank the Center for Advanced Research Computing (CARC) at the University of Southern California for providing the computational resources for this project. We also thank Mattia Polimanti at Sapienza Università di Roma for testing the reproducibility of our computational pipelines.

## SUPPLEMENTARY MATERIAL

The AncestryGrapher toolkit AppNote Supplementary Data.pdf contains the following: 1) Table S1 – Output file containing global ancestry information generated by `RFMix v.2`; 2) Figure S1 – Inferred global ancestry proportions for selected individuals from the Mozabite Berber population; 3) Figure S2 – Inferred global ancestry proportions for all individuals in the Mozabite Berber population from the HGDP-CEPH dataset; and 4) Figure S3 – Examples of local ancestry along chromosome 2 in the Mozabite Berber population.

## REFERENCES

1. Alexander DH, Novembre J, Lange K. Fast model-based estimation of ancestry in unrelated individuals. Genome Res 2009;19:1655–64.

2. Anguita-Ruiz A, Aguilera CM, Gil Á. Genetics of lactose intolerance: An updated review and online interactive world maps of phenotype and genotype frequencies. Nutrients 2020;12:2689.

3. Arauna LR, Mendoza-Revilla J, Mas-Sandoval A et al. Recent historical migrations have shaped the gene pool of Arabs and Berbers in North Africa. Mol Biol Evol 2016:msw218.

4. Bekada A, Arauna LR, Deba T et al. Genetic heterogeneity in Algerian human populations. PLoS One 2015;10:e0138453.

5. Bersaglieri T, Sabeti PC, Patterson N et al. Genetic signatures of strong recent positive selection at the lactase gene. Am J Hum Genet 2004;74:1111–20.

6. Browning SR, Browning BL, Zhou Y et al. Analysis of human sequence data reveals two pulses of archaic Denisovan admixture. Cell 2018;173:53-61.e9.

7. Browning SR, Waples RK, Browning BL. Fast, accurate local ancestry inference with FLARE. Am J Hum Genet 2023;110:326–35.

8. Campbell MC, Hirbo JB, Townsend JP et al. The peopling of the African continent and the diaspora into the new world. Curr Opin Genet Dev 2014;29:120–32.

9. Campbell MC, Ranciaro A. Human adaptation, demography and cattle domestication: an overview of the complexity of lactase persistence in Africa. Hum Mol Genet 2021;30:R98–109.

10. Campbell MC, Tishkoff SA. African genetic diversity: implications for human demographic history, modern human origins, and complex disease mapping. Annu Rev Genomics Hum Genet 2008;9:403–33.

11. Campbell MC, Tishkoff SA. The evolution of human genetic and phenotypic variation in Africa. Curr Biol 2010;20:R166–73.

12. Carrot-Zhang J, Han S, Zhou W et al. Analytical protocol to identify local ancestry-associated molecular features in cancer. STAR Protoc 2021;2:100766.

13. Cheng C-Y, Kao WHL, Patterson N et al. Admixture mapping of 15,280 African Americans identifies obesity susceptibility loci on chromosomes 5 and X. PLoS Genet 2009;5:e1000490.

14. Daya M, van der Merwe L, Gignoux CR et al. Using multi-way admixture mapping to elucidate TB susceptibility in the South African Coloured population. BMC Genomics 2014;15:1021.

15. Dias-Alves T, Mairal J, Blum MGB. Loter: A software package to infer local ancestry for a wide range of species. Mol Biol Evol 2018;35:2318–26.

16. Durand EY, Do CB, Mountain JL et al. Ancestry Composition: A novel, efficient pipeline for ancestry deconvolution. bioRxiv 2014, DOI: 10.1101/010512.

17. Enattah NS, Jensen TGK, Nielsen M et al. Independent introduction of two lactase-persistence alleles into human populations reflects different history of adaptation to milk culture. Am J Hum Genet 2008;82:57–72.

18. Enattah NS, Sahi T, Savilahti E et al. Identification of a variant associated with adult-type hypolactasia. Nat Genet 2002;30:233–7.

19. Freedman ML, Haiman CA, Patterson N et al. Admixture mapping identifies 8q24 as a prostate cancer risk locus in African-American men. Proc Natl Acad Sci U S A 2006;103:14068–73.

20. Gerbault P, Moret C, Currat M et al. Impact of selection and demography on the diffusion of lactase persistence. PLoS One 2009;4:e6369.

21. Gopalan S, Smith SP, Korunes K et al. Human genetic admixture through the lens of population genomics. Philos Trans R Soc Lond B Biol Sci 2022;377:20200410.

22. Guan Y. Detecting structure of haplotypes and local ancestry. Genetics 2014;196:625–42.

23. Gurdasani D, Carstensen T, Tekola-Ayele F et al. The African Genome Variation Project shapes medical genetics in Africa. Nature 2015;517:327–32.

24. Hassan HY, van Erp A, Jaeger M et al. Genetic diversity of lactase persistence in East African populations. BMC Res Notes 2016;9:8.

25. Hellenthal G, Busby GBJ, Band G et al. A genetic atlas of human admixture history. Science 2014;343:747–51.

26. Henn BM, Botigué LR, Gravel S et al. Genomic ancestry of North Africans supports back-to-Africa migrations. PLoS Genet 2012;8:e1002397.

27. Hilmarsson H, Kumar AS, Rastogi R et al. High resolution ancestry deconvolution for next generation genomic data. bioRxiv 2021, DOI: 10.1101/2021.09.19.460980.

28. Hollfelder N, Babiker H, Granehäll L et al. The genetic variation of lactase persistence alleles in northeast Africa. bioRxiv 2020, DOI: 10.1101/2020.04.23.057356.

29. Ingram CJE, Elamin MF, Mulcare CA et al. A novel polymorphism associated with lactose tolerance in Africa: multiple causes for lactase persistence? Hum Genet 2007;120:779–88.

30. Ingram CJE, Raga TO, Tarekegn A et al. Multiple rare variants as a cause of a common phenotype: several different lactase persistence associated alleles in a single ethnic group. J Mol Evol 2009;69:579–88.

31. Itan Y, Powell A, Beaumont MA et al. The origins of lactase persistence in Europe. PLoS Comput Biol 2009;5:e1000491.

32. Jones BL, Raga TO, Liebert A et al. Diversity of lactase persistence alleles in Ethiopia: signature of a soft selective sweep. Am J Hum Genet 2013;93:538–44.

33. Koller D, Wendt FR, Pathak GA et al. Denisovan and Neanderthal archaic introgression differentially impacted the genetics of complex traits in modern populations. BMC Biol 2022;20:249.

34. Korunes KL, Goldberg A. Human genetic admixture. PLoS Genet 2021;17:e1009374.

35. Macholdt E, Lede V, Barbieri C et al. Tracing pastoralist migrations to southern Africa with lactase persistence alleles. Curr Biol 2014;24:875–9.

36. Maples BK, Gravel S, Kenny EE et al. RFMix: a discriminative modeling approach for rapid and robust local-ancestry inference. Am J Hum Genet 2013;93:278–88.

37. Montserrat DM, Bustamante C, Ioannidis A. Lai-net: Local-ancestry inference with neural networks. ICASSP 2020 - 2020 IEEE International Conference on Acoustics, Speech and Signal Processing (ICASSP). IEEE, 2020.

38. Price AL, Tandon A, Patterson N et al. Sensitive detection of chromosomal segments of distinct ancestry in admixed populations. PLoS Genet 2009;5:e1000519.

39. Pritchard JK, Stephens M, Donnelly P. Inference of population structure using multilocus genotype data. Genetics 2000;155:945–59.

40. Qin P, Stoneking M. Denisovan ancestry in east Eurasian and Native American populations. Mol Biol Evol 2015;32:2665–74.

41. Ranciaro A, Campbell MC, Hirbo JB et al. Genetic origins of lactase persistence and the spread of pastoralism in Africa. Am J Hum Genet 2014;94:496–510.

42. Rudan I. Health effects of human population isolation and admixture. Croat Med J 2006;47:526–31.

43. Sánchez-Quinto F, Lalueza-Fox C. Almost 20 years of Neanderthal palaeogenetics: adaptation, admixture, diversity, demography and extinction. Philos Trans R Soc Lond B Biol Sci 2015;370:20130374.

44. Sankararaman S, Mallick S, Patterson N et al. The combined landscape of Denisovan and neanderthal ancestry in present-day humans. Curr Biol 2016;26:1241–7.

45. Sankararaman S, Sridhar S, Kimmel G et al. Estimating local ancestry in admixed populations. Am J Hum Genet 2008;82:290–303.

46. Scheinfeldt LB, Soi S, Lambert C et al. Genomic evidence for shared common ancestry of East African hunting-gathering populations and insights into local adaptation. Proc Natl Acad Sci U S A 2019;116:4166–75.

47. Silva M, Oteo-García G, Martiniano R et al. Biomolecular insights into North African-related ancestry, mobility and diet in eleventh-century Al-Andalus. Sci Rep 2021;11:18121.

48. Sundquist A, Fratkin E, Do CB et al. Effect of genetic divergence in identifying ancestral origin using HAPAA. Genome Res 2008;18:676–82.

49. Tang H, Coram M, Wang P et al. Reconstructing genetic ancestry blocks in admixed individuals. Am J Hum Genet 2006;79:1–12.

50. The 1000 Genomes Project Consortium. An integrated map of genetic variation from 1,092 human genomes. Nature 2012;491:56–65.

51. Tishkoff SA, Reed FA, Ranciaro A et al. Convergent adaptation of human lactase persistence in Africa and Europe. Nat Genet 2007;39:31–40.

52. Uren C, Hoal EG, Möller M. Putting RFMix and ADMIXTURE to the test in a complex admixed population. BMC Genet 2020;21:40.

53. Uren C, Kim M, Martin AR et al. Fine-scale human population structure in southern Africa reflects ecogeographic boundaries. Genetics 2016;204:303–14.

54. Vespasiani DM, Jacobs GS, Cook LE et al. Denisovan introgression has shaped the immune system of present-day Papuans. PLoS Genet 2022;18:e1010470.

55. Vicente M, Priehodová E, Diallo I et al. Population history and genetic adaptation of the Fulani nomads: inferences from genome-wide data and the lactase persistence trait. BMC Genomics 2019;20:915.

56. Vyas DN, Mulligan CJ. Analyses of Neanderthal introgression suggest that Levantine and southern Arabian populations have a shared population history. Am J Phys Anthropol 2019;169:227–39.

57. Wall JD, Yoshihara Caldeira Brandt D. Archaic admixture in human history. Curr Opin Genet Dev 2016;41:93–7.

58. Witt KE, Funk A, Añorve-Garibay V et al. The impact of modern admixture on archaic human ancestry in human populations. Genome Biol Evol 2023;15, DOI: 10.1093/gbe/evad066.

