## Supplementary Data for "AncestryGrapher toolkit: Python command-line pipelines to visualize global– and local– ancestry inferences from the RFMix2 software"

**Table S1—Output file containing global ancestry information generated by RFMix v.2**. The global ancestry output files contain tab-separated content that corresponds to the following: name of recipient and donor population (column 1); proportion of ancestry (column 2, column 3, and column 4) for a given chromosome. The order and names of the reference populations used in ancestry determination are also given in the header row.

```
#rfmix diploid global ancestry .Q format output
#sample	Africa	Europe	Middle_East
Mozabite1	0.13847	0.22469	0.63684
Yoruba1	1.00000	0.00000	0.00000
Yoruba2	1.00000	0.00000	0.00000
Yoruba3	1.00000	0.00000	0.00000
Yoruba4	1.00000	0.00000	0.00000
Yoruba5	1.00000	0.00000	0.00000
Yoruba6	1.00000	0.00000	0.00000
Yoruba7	1.00000	0.00000	0.00000
Yoruba8	1.00000	0.00000	0.00000
Yoruba9	1.00000	0.00000	0.00000
Yoruba10	1.00000	0.00000	0.00000
Yoruba11	1.00000	0.00000	0.00000
Yoruba12	1.00000	0.00000	0.00000
Yoruba13	1.00000	0.00000	0.00000
Yoruba14	1.00000	0.00000	0.00000
Yoruba15	1.00000	0.00000	0.00000
Yoruba16	1.00000	0.00000	0.00000
Yoruba17	1.00000	0.00000	0.00000
Yoruba18	1.00000	0.00000	0.00000
Yoruba19	1.00000	0.00000	0.00000
Yoruba20	1.00000	0.00000	0.00000
Yoruba21	1.00000	0.00000	0.00000
Yoruba22	1.00000	0.00000	0.00000
```

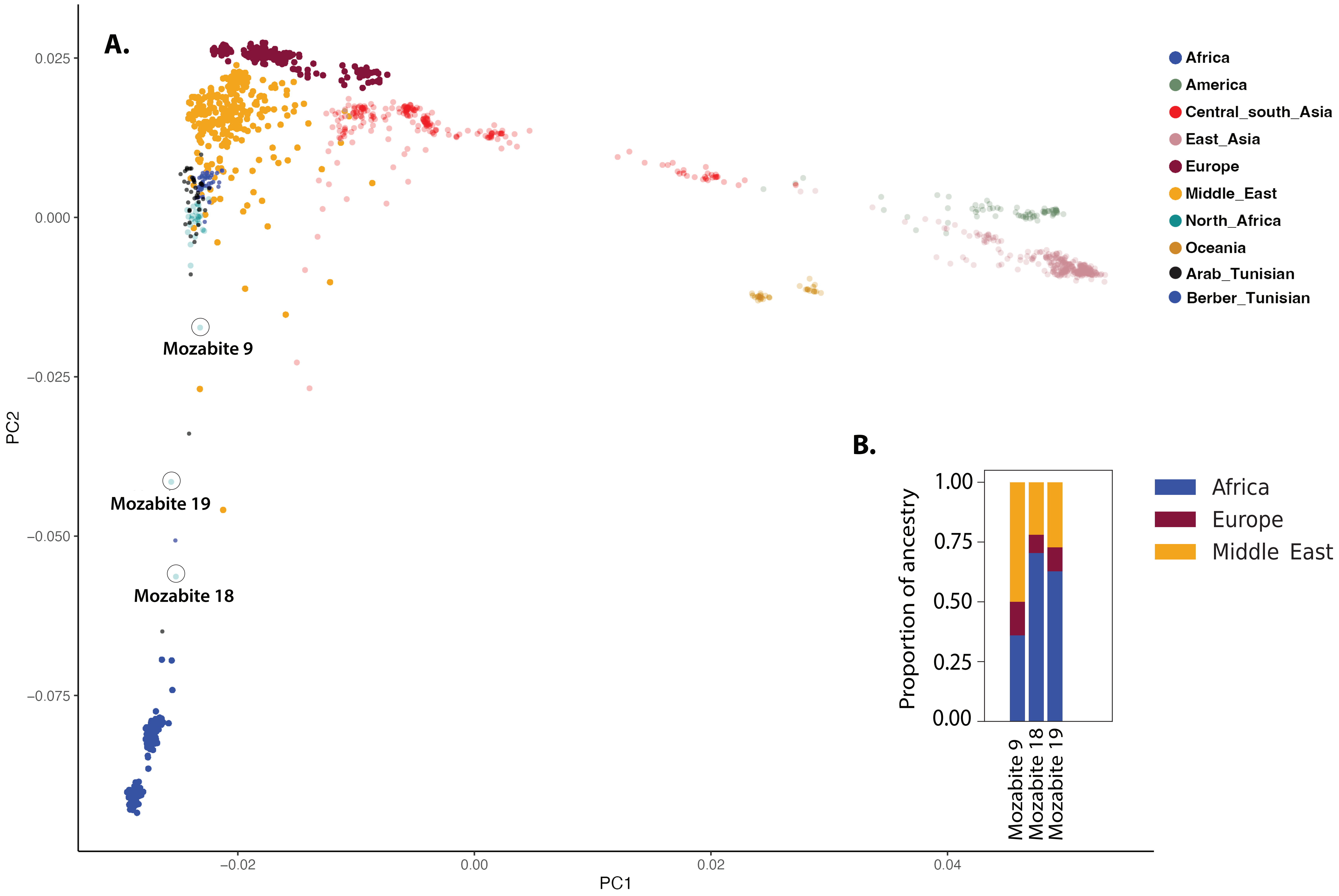


**Figure S1: Inferred global ancestry proportions for selected individuals from the Mozabite Berber population**. To explore genetic structure in populations, we performed a Principal Component Analysis (PCA) of global populations. In this plot, each dot represents an individual and is colored based on geographic origin (e.g., the Americas, Central/South Asia, East Asia, Europe, the Middle East, and Oceania). PC1 and PC2 are given on the x-axis and y-axis, respectively. In addition, we plotted the output from RFMix v. 2 using the GAP algorithm to visualize the global ancestry in selected individuals from the Mozabite Berber population which are circled in the PCA (Panel B). In this plot, each color represents a distinct inferred ancestry and the reference populations used in this ancestry determination are given in the figure legend.

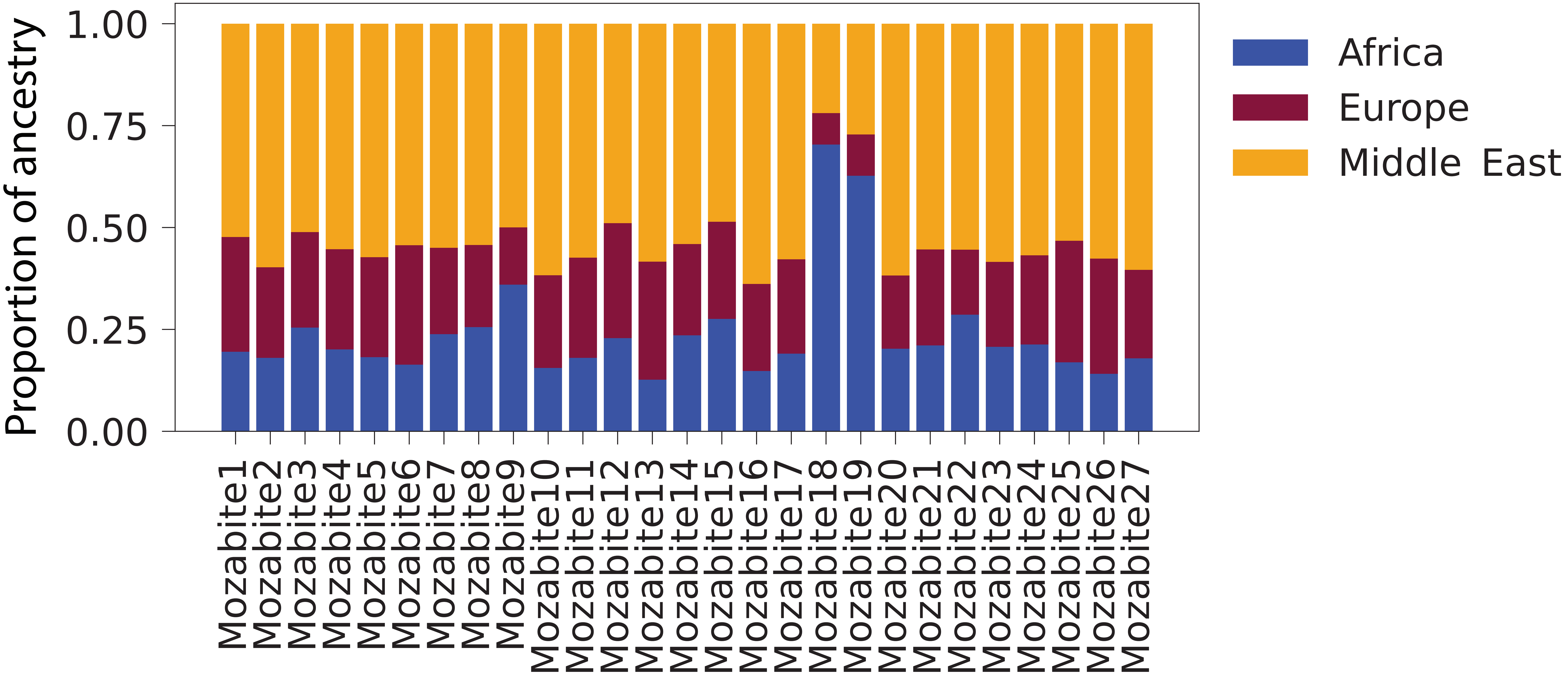


**Figure S2: Inferred global ancestry proportions for all individuals in the Mozabite Berber population from the HGDP-CEPH dataset**. We plotted the output from RFMix v.2 using the GAP algorithm to visualize the global ancestry of individuals from the Mozabite Berber population. In this plot, each color represents a distinct inferred ancestry and the reference populations used in this ancestry determination are given in the figure legend.

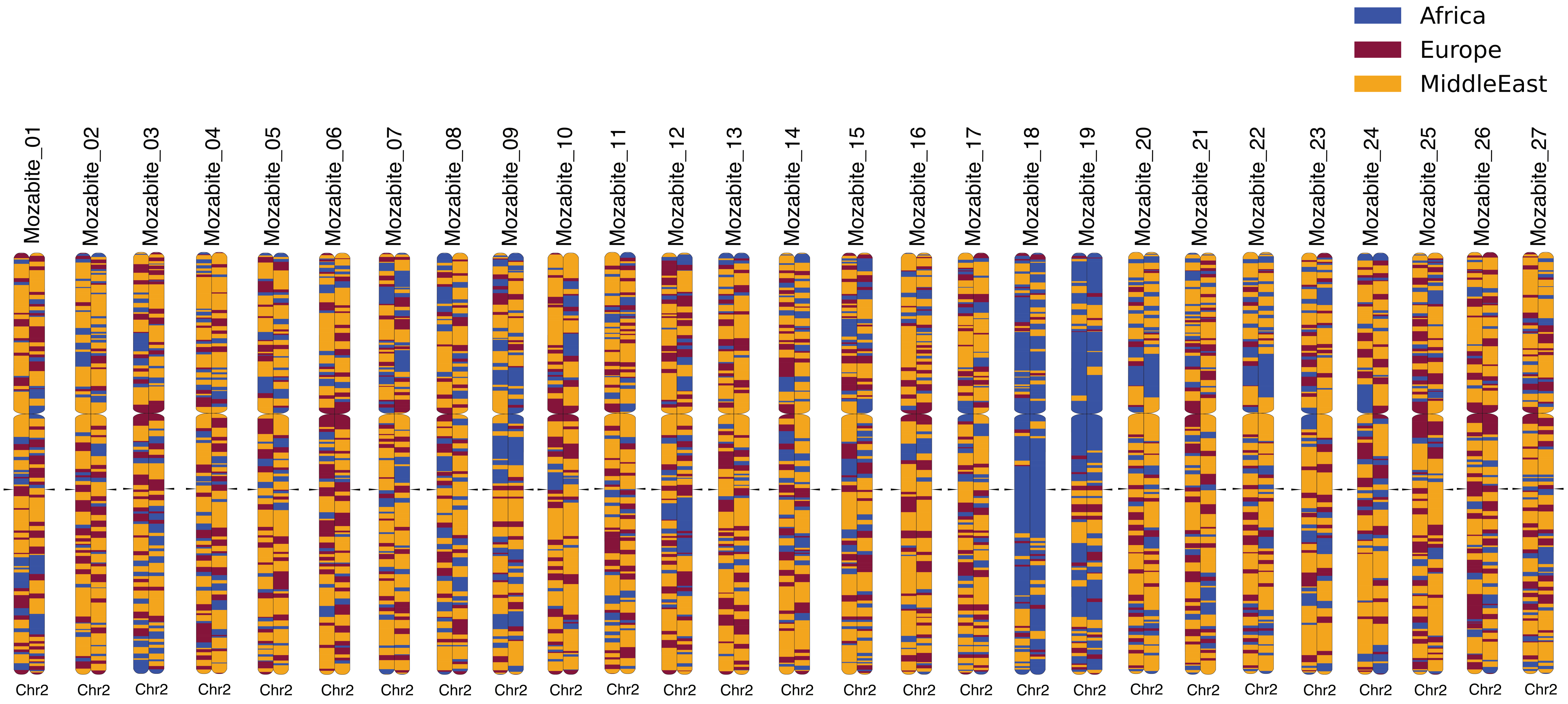


**Figure S3: Examples of local ancestry along chromosome 2 in the Mozabite Berber population**. These plots show diploid chromosomes for each individual in the Mozabite population from the publicly available HGDP-CEPH dataset. The orange color indicates Middle Eastern ancestry; purple represents European ancestry; and blue signifies ancestry originating from sub-Saharan Africa. The arrows pointing toward chromosome 2 indicate the chromosomal region containing common alleles associated with lactase persistence.
